## Supplementary figures and images for "A microRNA program controls the transition of cardiomyocyte hyperplasia to hypertrophy and stimulates mammalian cardiac regeneration"

### Supplemental Figure 1

#### Supplemental Figure 1

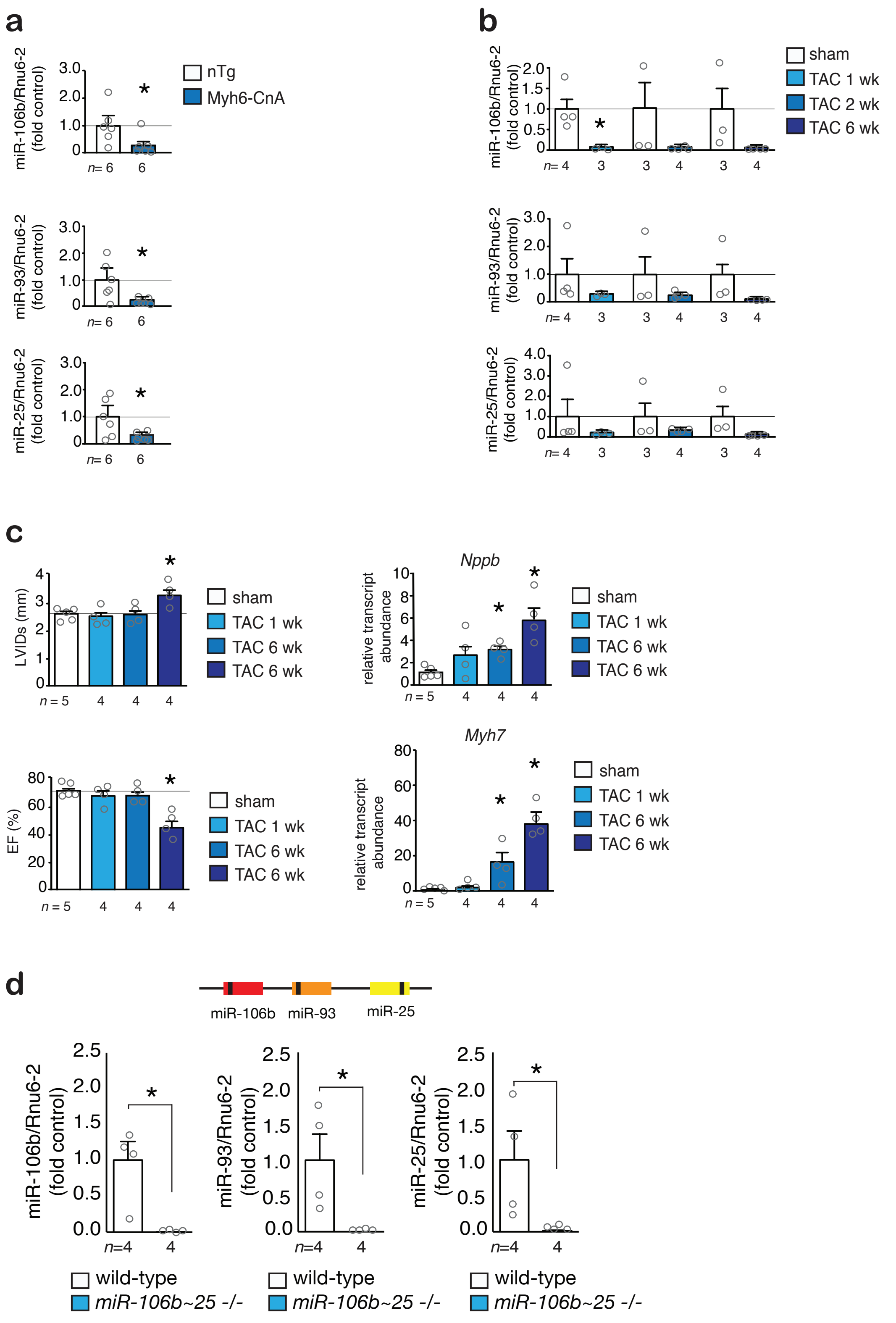

### Supplemental Figure 2

#### Supplemental Figure 2

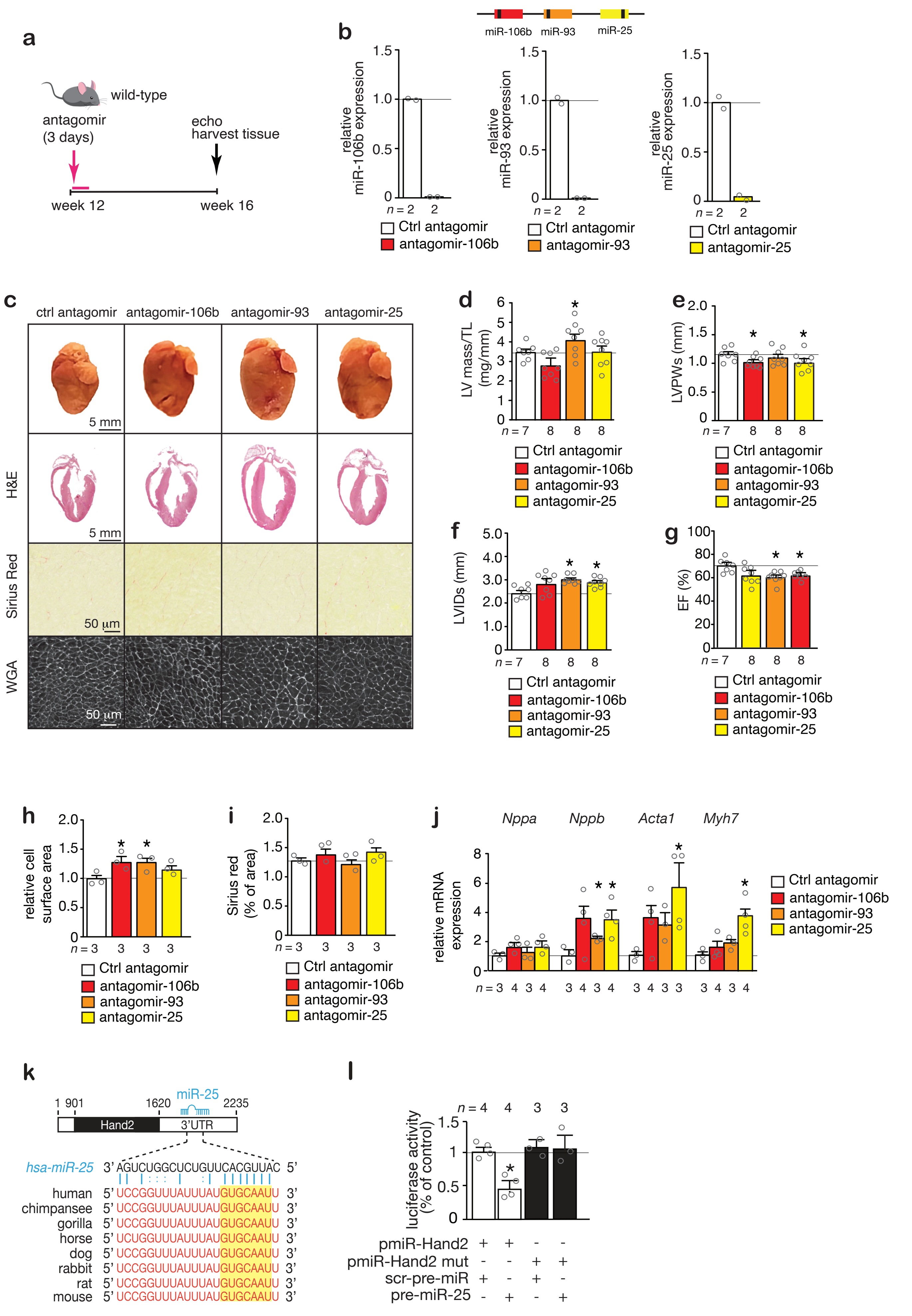

### Supplemental Figure 3

#### Supplemental Figure 3

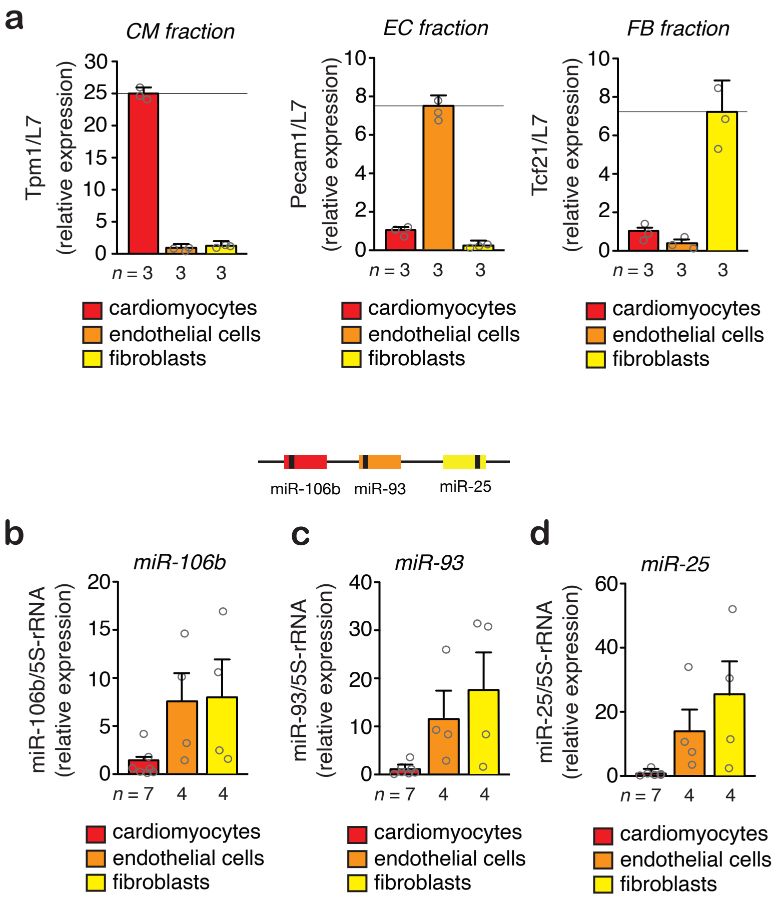

### Supplemental Figure 5

#### Supplemental Figure 5

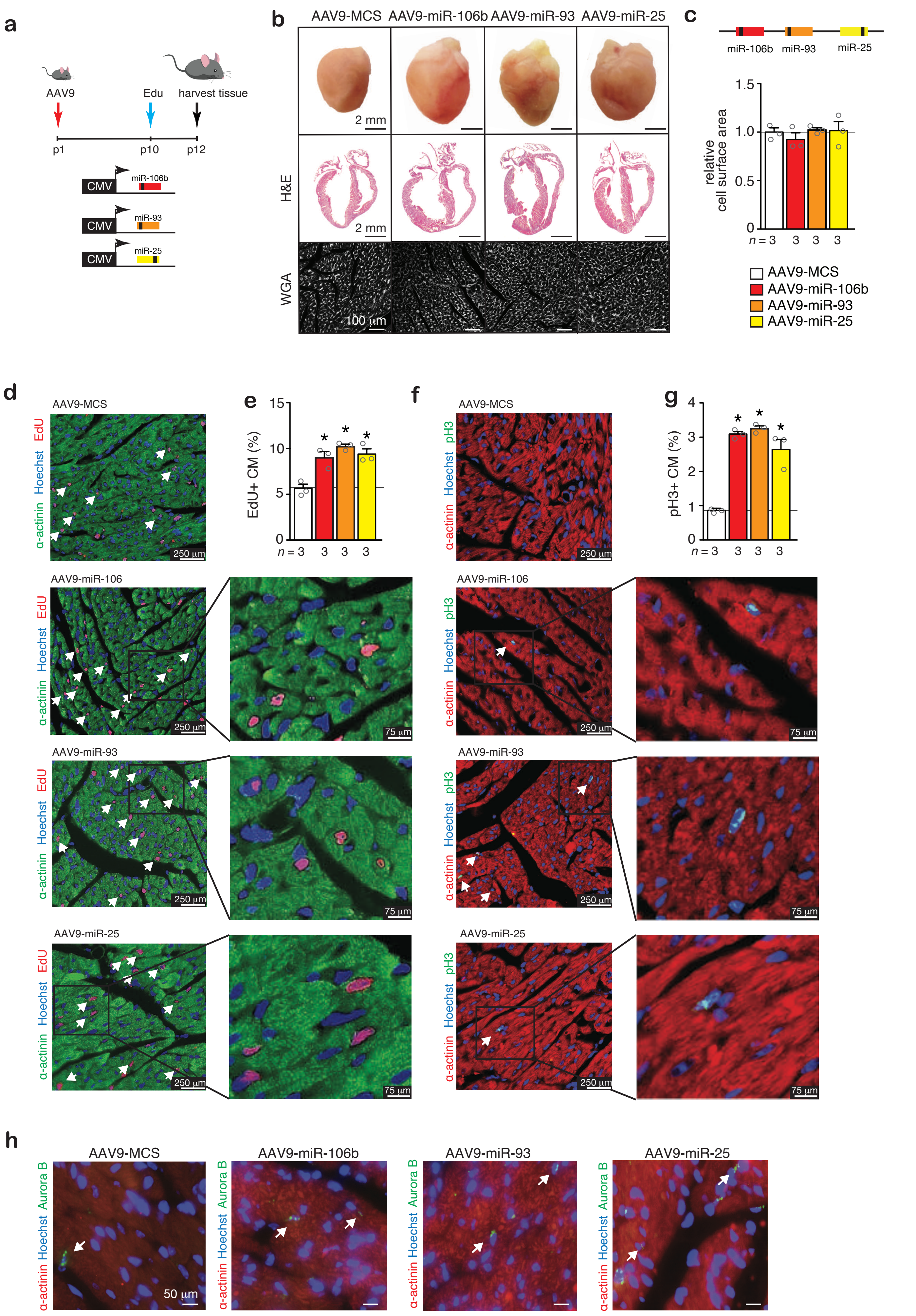

### Supplemental Figure 6

#### Supplemental Figure 6

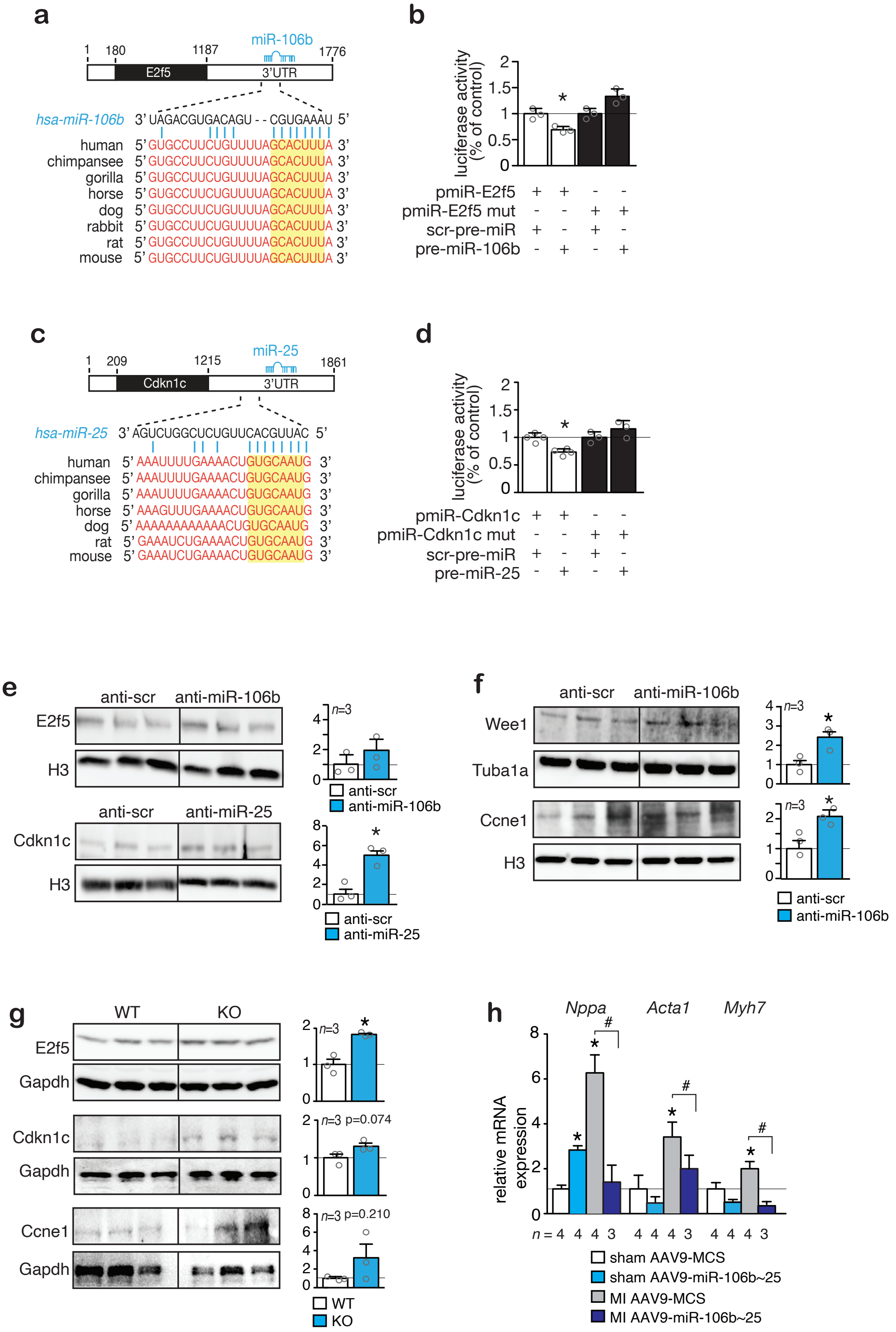
