## Supplemental Figure 4 for "A microRNA program controls the transition of cardiomyocyte hyperplasia to hypertrophy and stimulates mammalian cardiac regeneration"

**a**

**AAV9-MCS**

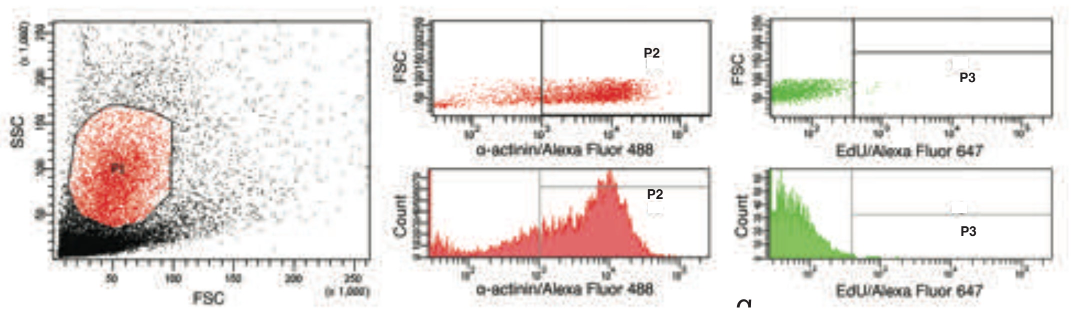


| Population | #Events | %Parent | %Total |
|---|---|---|---|
| All Events | 20,000 | 100.0 | 100.0 |
| P1_Analyzed Events | 4,879 | 22.2 | 22.2 |
| P2_α-actinin+CMs | 3,543 | 72.6 | 16.1 |
| P3_EdU+CMs | 20 | 0.6 | 0.1 |

**AAV9-miR-106~25**

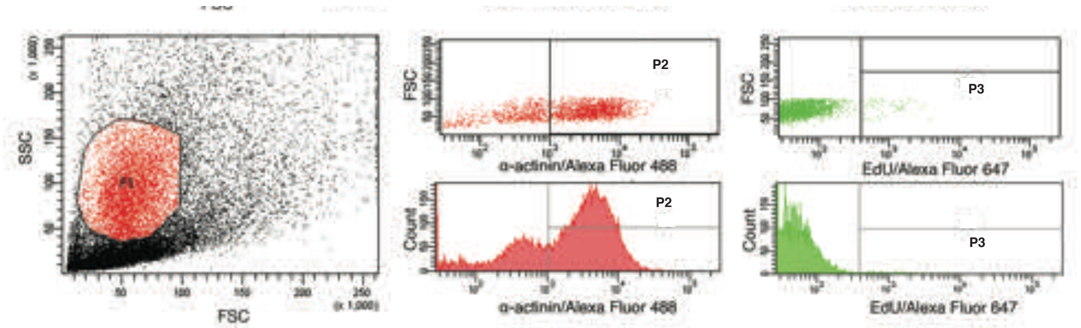


| Population | #Events | %Parent | %Total |
|---|---|---|---|
| All Events | 60,000 | 100.0 | 100.0 |
| P1_Analyzed Events | 11,473 | 19.1 | 19.1 |
| P2_α-actinin+CMs | 8,041 | 70.1 | 13.4 |
| P3_EdU+CMs | 101 | 1.3 | 0.2 |

**b**

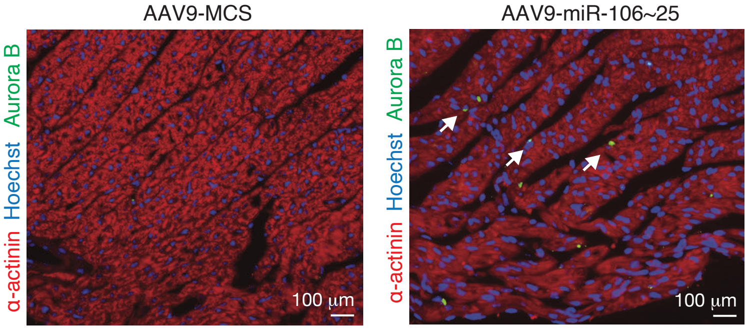
